# Heliozoan cellular architecture predicts prey capture performance and growth rate

**DOI:** 10.64898/2026.10.19.760065

**Authors:** Paul E Schavemaker, Michael Lynch

**Affiliations:** Biodesign Center for Mechanisms of Evolution, Arizona State University, Tempe, AZ, 85287, USA; Living Systems Institute, University of Exeter, Stocker Road, Exeter, EX4 4QD, UK

## Abstract

Prey capture is essential for growth in many species of single-celled eukaryotes, and its various elaborations phenotypically define major eukaryotic clades. Despite impressive work on the mechanistic basis, capture rates, and trade-offs in prey capture, in flagellates in particular, no quantitative assessment of the full fitness contributions of specific prey capture mechanisms has been performed. Here, a step in the direction of such an assessment is taken for the axopod-based prey capture mechanism of heliozoans. A model was established that (1) obtains prey capture rates from a geometrically accurate simulation of prey capture, (2) assesses the energetic cost of the prey capture mechanism as a function of axopod length and number, and (3) uses the capture rates and cost to calculate a cell division time. The heliozoan prey capture model captures the essence of the mechanism and performs decent predictions of axopod cost, axopod length, axopod number, prey capture rate, clearance rate, and cell division time. The core elements of the capture model are broadly applicable and, when supported by mechanistic information, can be applied to flagellates, ciliates, amoeba and other organisms, forming a basis for elucidating how such varied mechanisms of prey capture were able to evolve.

## Introduction

Examining prey capture mechanisms among single-celled eukaryotes (protists) leaves one with a sense of bewilderment. Ciliates, amoebae, choanoflagellates, dinoflagellates, and heliozoans all perform essentially the same activity, capturing prey, but do so by a staggering variety of mechanisms. The relation between the specific prey capture mechanisms and the character of the prey or environment remains largely a mystery. This is unfortunate because these prey capture mechanisms are responsible for much of the stunning morphological diversity present in single-celled eukaryotes and are likely to have had central roles in establishing the cellular *Bauplans* of large clades of protist life.

It has been hypothesized that the last eukaryotic common ancestor was a phagotrophic flagellate^1,2^ perhaps similar in phenotype to the present-day excavates^3,4^. Some of these excavates capture prey cells by beating a vaned flagellum in a groove to generate a feeding current and engulfing the prey by phagocytosis^3^. Mechanistic studies of prey capture^5,6^ are of considerable evolutionary interest because they investigate the effect of presence or absence of traits on prey capture ability. For example, in the excavates it was demonstrated that the presence of a feeding groove and a flagellar vane is important for prey capture^3^. In dinoflagellates, the presence or absence of the transverse and longitudinal flagellum, as well as the presence or absence of flagellar hairs, was found to affect the prey clearance rate in mathematical models of flow^7^. Flagella also work together with other cellular appendages to facilitate prey capture, for example the haptonema in haptophytes^8–10^ and collar microvilli in choanoflagellates^11,12^.

The flagellated protists, diverse though they are^13^, are only one of several phenotypically identifiable groups with distinct prey capture mechanisms. Amoebae have the distinguishing characteristic of deforming their entire cell body to move and capture prey^14–20^, and include strange forms such as vampyrellids^21,22^. Ciliates encompass multiple distinct predatory styles^23^ for example the rapid extensions and contractions of the neck of *Lacrymaria*^24,25^, the capture and engulfment of prey equal in size to the predator by *Didinium*^26–28^, as well as various other forms^29–34^. The suctorians are a clade within the ciliates that lack flagella in their vegetative state and capture prey by tentacles^35–37^. Other mechanisms of prey capture are by a reticulopodium as in foraminifera^38^ and by a single large tentacle as in the atypical dinoflagellate *Noctiluca scintillans*^39,40^.

Heliozoa, the focus of this work, are a phenotypically defined grouping of single-celled eukaryotes that have spherical cell bodies festooned with straight appendages, called axopods, that emerge from the cell surface at right angles and are used for prey capture^41^. Heliozoa occur in the water column^41–43^ as well as the benthos^41^ (see also *Actinosphaerium* in Esteban and Fenchel^44^, p. 137). Axopod-based prey capture and phagocytosis were observed for *Actinophrys sol*^45^ already in 1848. Heliozoa typically aren’t motile, so prey is captured passively^41^. As prey collides with an axopod it gets stuck by an adhesive that is probably excreted by extrusomes^46,47^. Kinetocysts that connect predator axopod to prey have been observed in *Raphidiophrys contractilis*^48^. After capture, the prey is transported to the cell surface either by contraction of the axopod or by surface transport along the axopod^47–49^. Heliozoa feed on a range of prey of different sizes: bacteria, algae, ciliates, and rotifers^41,42,50–52^. *Raphidocystis contractilis* cells can collaborate to capture and digest the larger *Actinophrys sol* by utilizing a shared vacuole^53^.

Prey capture mechanisms are shaped by their ability to improve fitness. Thus, the persistence of these traits and their occasional reappearance in independent lineages (convergent evolution) requires an explanation grounded in an assessment of their fitness contributions. In this assessment the parameters that govern the prey capture mechanism must be varied systematically. For example, for an excavate-like flagellate one would have to vary the length of the flagellum and the depth of the feeding groove (Figure 1A), for choanoflagellates one would have to vary collar microvilli length and number, for amoebae one would have to vary the enclosed volume and thickness of the phagocytic cup.

**Figure 1:**
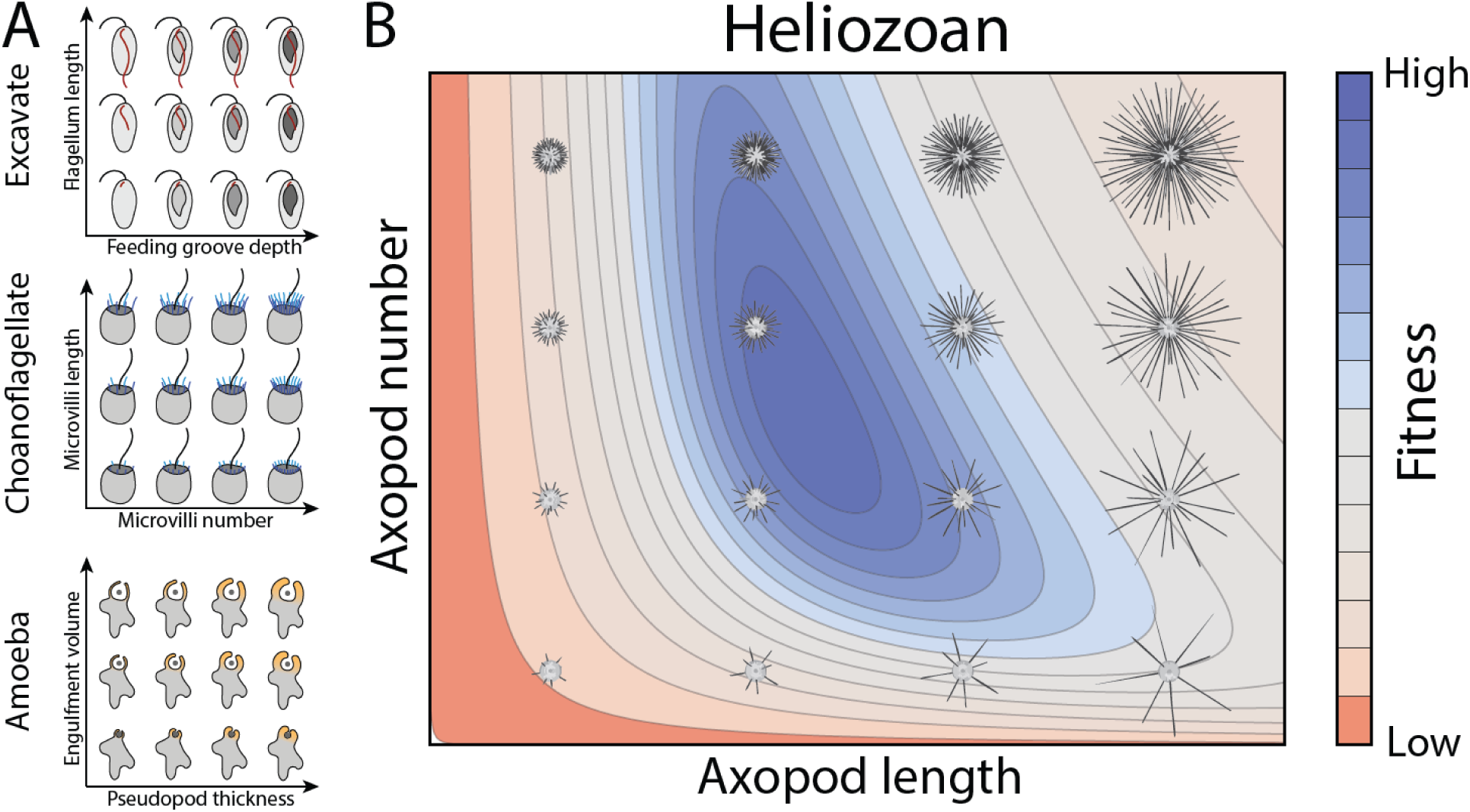
Fitness landscapes for prey capture mechanisms. A) Potential ways in which key traits could affect the fitness contribution of prey capture mechanisms. B) An imagined fitness landscape for the heliozoan prey capture mechanism that depends on axopod number and length.

Partial assessments of fitness have previously been performed for prey capture mechanisms^3,7^. However, a calculation of fitness, as opposed to the improvement of a single cellular activity, in which both costs and gains are assessed and integrated, and in which the key parameters that govern the character of the trait are varied systematically, has not been undertaken. This work provides such an assessment on a simplified model of heliozoan axopod-based prey capture. The choice for the relatively rare heliozoan phenotype as opposed to, say, flagellates, is the relative straightforward nature of the prey capture mechanism. Demonstrating these fitness calculations for a (relatively speaking) simple prey capture mechanism will spur these developments for other mechanisms, such as flagella- or pseudopod-based prey capture. The heliozoan mechanism is of interest also because it has convergently evolved multiple times and has persisted for a long time and is thus not idiosyncratic.

## Methods

### Calculating energetic cost of heliozoan prey capture

Heliozoa capture prey with axopods, which cost resources to produce. Here, the cost of axopods is expressed in units of ATP, in line with earlier work^54–60^. Axopods are composed of the following main constituents (Figure 2A): a membrane that envelops the axopod, microtubules that extend from the cell surface outward and that shape the axopod, and the extensions of the axopod microtubules to the cell center where they combine into a structure called the centroplast^61^ or axoplast^62^. The calculation of the axopod costs is identical to what has been described previously^58^ (Supplementary Information). The cost of the prey capture apparatus, *C_cap_*, is the sum of the cost of all axopods (Eq. S1-S4):

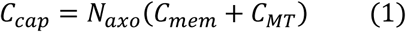

**Figure 2:**
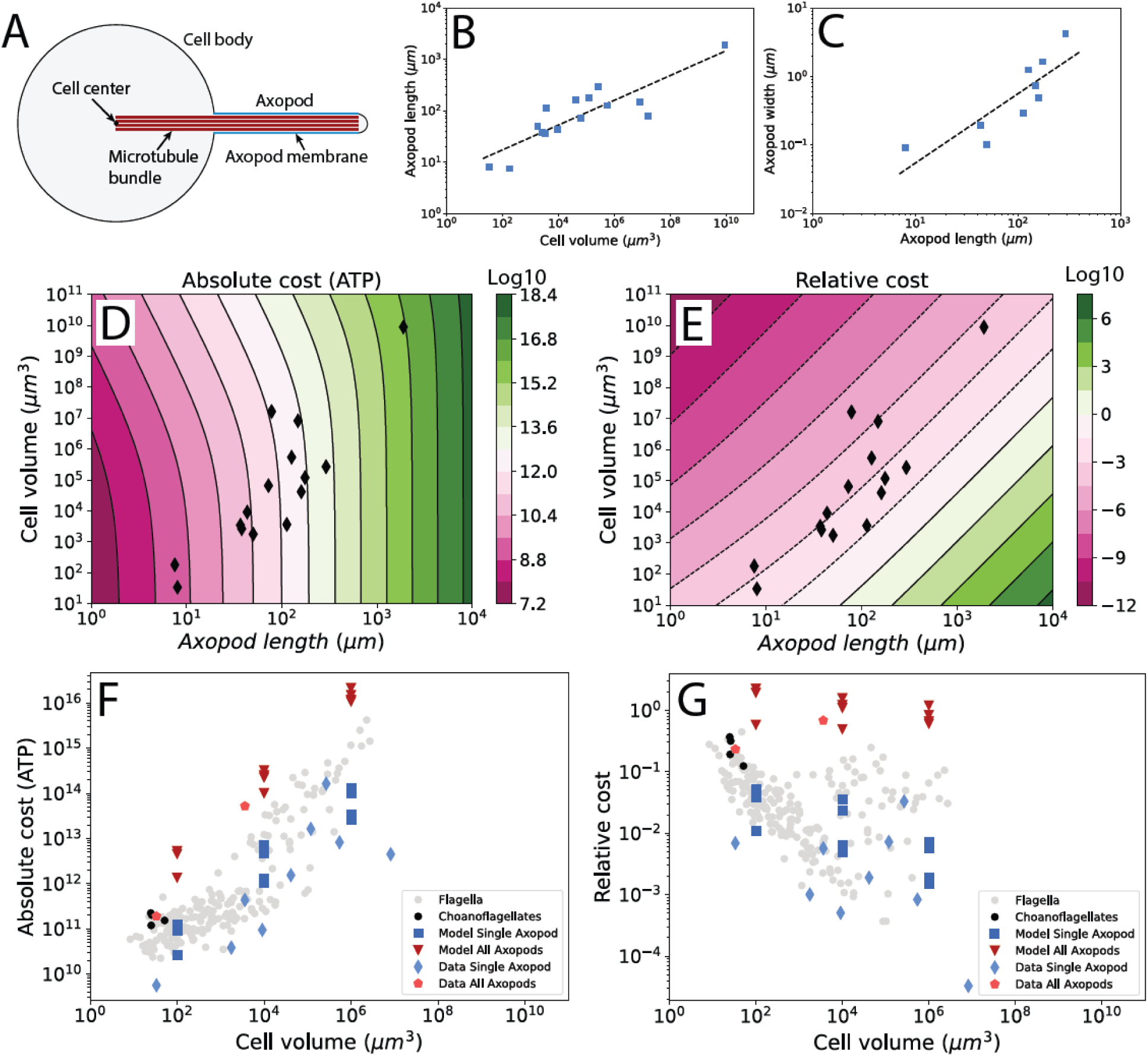
Quantitation of axopod dimensions and construction costs in Heliozoa. A) Axopod features included in cost calculations. B) Axopod length and cell volume for diverse species of heliozoa (*y* = 5.7*x*^0.24^). C) Axopod width and axopod length for diverse species of heliozoa (*y* = 0.0053*x*^1.01^). D) Absolute construction cost, in units of ATP hydrolyses (including opportunity costs), of individual axopods (Eq. S1-S3). Black diamonds: axopod costs for specific heliozoan species, assuming the scaling in C. E) Relative construction costs of axopods (absolute cost divided by the cost of the cell body). Black diamonds as in D. F) Absolute construction cost for single axopods or full complements of axopods, comparing data to optima obtained from the cell division time calculations. Construction costs of flagella (full complement, not individual flagella) across eukaryotic diversity (including ciliates and flagellates)^56,57,60^ and of choanoflagellate prey capture (flagella + microvilli). G) Same as F but for relative construction costs.

Here, *N_axo_* is the number of axopods on a single heliozoan cell, *C_mem_* is the cost of the membrane, and *C_MT_* is the cost of the microtubules.

### Modeling the rate of capturing diffusing prey

The prey capture rate is determined as a function of axopod number, axopod length and prey concentration for two types of prey: (1) prey that moves by diffusion and (2) prey that swims. The rate of capturing diffusing prey is calculated using the reaction-diffusion software Smoldyn^58,63^ (Supplementary Information). This simulation includes a detailed heliozoan geometry with a sphere representing the cell body and cylinders representing the axopods. Prey cells diffuse, and when they collide with the axopods, they are captured. Prey capture rates are determined for different axopod lengths (*L_axo_*), axopod numbers (*N_axo_*), and predator volumes. For use in cell division time calculations, the simulation output is fitted with:

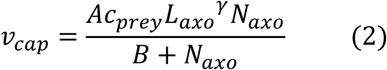

Here, *c_prey_* is prey concentration and *A*, *B*, and *γ* are fit parameters.

### Modeling the rate of capturing swimming prey

The rate of capturing swimming prey is calculated from a combination of two simulations written in Python (Figure 3A, Supplementary Information). The first simulation determines the rate at which swimming prey collide with a boundary sphere, positioned concentrically with the heliozoan predator, that has its outer boundary at the tips of the axopods (Figure 3B). The second simulation determines the probability that a swimming prey cell, located within the boundary sphere, will collide with an axopod. Combined, these two simulations give the prey capture rate (Equation 2), which is determined for different axopod lengths, axopod numbers, predator volumes, and prey volumes.

**Figure 3:**
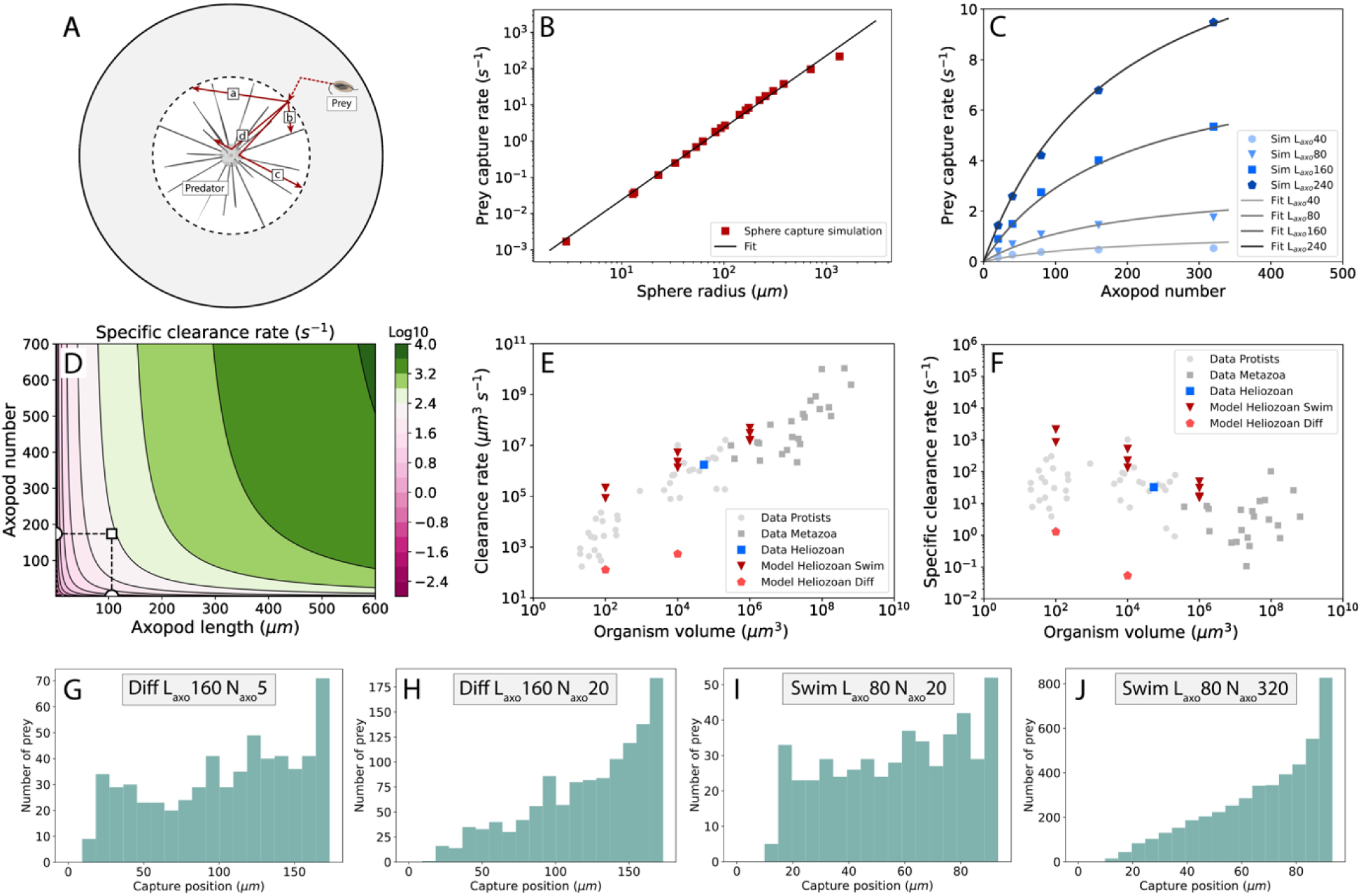
Quantification of heliozoan prey capture rate. A) Model of prey capture of swimming prey by heliozoa. Labels: “a” escape, “b” direct capture, “c” indirect escape, and “d” indirect capture. The grey area between the solid and dashed circle is simulated in the first step, and the area within the dashed circle is simulated in the second step. B) Capture rate by a sphere as a function of its radius (dashed circle in A). First step of the simulation. Prey concentration: 10^-6^ µm^-3^, regression: *y* = 2.5 × 10^−4^*x*^2^. C) Swimming prey capture rate from the heliozoan prey capture model for different axopod numbers and lengths. Prey concentration: 10^-6^ µm^-3^. A fit with Eq. 2 gives *A* = 6546, *B* = 191, and *γ* = 1.41. D) Specific clearance rate calculated from the capture model. V_predator_: 10^4^ µm^3^, V_prey_: 10^2^ µm^3^. White square: position of minimal cell division time (Fig. 4B). E) Clearance rates calculated from optima in the prey capture models (diffusing and swimming prey) compared to experimental data^43,79^. F) Same as E but for specific clearance rate. G-J) Positions of prey capture along the axopod, with zero indicating the cell center. G-H: diffusing prey, and I-J: swimming prey. H and J have close to optimal axopod length and number. G and I have axopod numbers significantly lower than the optimum.

### Calculating the heliozoan cell division time

The heliozoan cell division time is calculated from a ratio of the nutrient requirement and the nutrient uptake rate. The nutrient requirement depends on the sum of the cost of the cell body (excluding axopods) and the cost of the axopods. As axopods are increased in number and length their costs also increase. The nutrient uptake rate depends on the axopod number and length as well, with the cell division time being decided by a balance between the two effects (Supplementary Information):

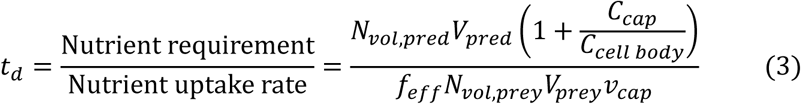

Here, *N_vol_*_,*pred*_ is the nutrient requirement for producing a unit of heliozoan cell volume, *N_vol_*_,*prey*_ is the number of nutrients present in the biomass of the prey cell, *V_pred_* and *V_prey_* are the cell volumes of the prey and predator (excluding the axopods), *C_cell body_* is the cost of producing an additional cell body in units of ATP, *f_eff_* is the conversion efficiency for converting between prey and predator biomass (here 0.25, Supplementary Information), *C_cap_* is the prey capture cost (Eq. 1), and *v_cap_* is the prey capture rate (Eq. 2).

### Modeling axopod saturation

Axopods don’t process prey instantaneously. This means that at high enough prey concentration, prey will collide with axopods that are already occupied with a previous catch, reducing the capture rate. An effective axopod number is introduced, *N_axo_*_,*eff*_, that depends on the processing time, *t_proc_*, and can account for the impact of high prey concentration (Supplementary Information, Eq. S24-S28). The modified prey capture rate is used with Eq. 3 to determine a saturation dependent cell division time.

### Dual prey model

The model presented in Eq. 3 is limited to one prey type at a time. To account for multiple prey types in the environment at the same time, Eq. 3 is modified to include two different prey cell volumes and two different prey capture rates (Supplementary Information, Eq. S31). In addition, the prey capture rates were adjusted to include a limit on the total volume of prey per unit volume of medium (Supplementary Information, Eq. S31). The fraction of the total prey volume that is smaller prey is given by *f_small_*.

## Results

### Quantitative cellular morphology of Heliozoans

Cellular morphology was examined quantitatively across heliozoan diversity using data collected from the literature^50,62,64–73^ (Table S1, Supplementary Information). The heliozoans in this dataset present spherical cell bodies with volumes ranging from 34 µm^3^ to 8.8×10^9^ µm^3^ (Figure 2B). The axopod lengths vary from 7.5 to 1892 µm (Figure 2B), and axopod widths from 0.092-4.2 µm (Figure 2C). Axopod numbers were collected from three species and ranged from 34-120. Density of microtubules in an axopod cross-section, required for axopod cost determinations, are 819±177 µm^-2^ (mean±SD, N_species_=3).

### Energetic cost of heliozoan prey capture

Cellular traits may present benefits for the cell, but typically also come with a construction-related resource cost. Resources come in various forms, elements^74–76^, amino acids, cell surface area occupancy^58,60,77,78^, etc.^60^, but are here expressed in units of energy, i.e. ATP, and include opportunity costs^54,60^. Heliozoan prey capture is carried out by axopods, and axopod costs include membrane and microtubule costs. We determine the cost of individual axopods as well as the prey capture mechanism as a whole (all axopods on the cell combined). These costs are determined for individual species of heliozoans as well as for the prey capture in the heliozoan prey capture models (Eq. 1-3). Both absolute costs, in units of ATP, and costs relative to the whole cell body were determined.

The absolute cost of individual axopods as a function of axopod length and cell volume (excluding axopods) is shown in Figure 2D. This reveals a strong relation between the cost of an axopod and the cell volume of the heliozoan (black diamonds), caused by an increase in axopod length with cell volume. The axopod length varies by only 250-fold, but the cost varies by about seven orders of magnitude. This is explained by the linear relation between axopod length and axopod width (Figure 2C) causing the cost of the axopod to scale with axopod length to the third power. Absolute axopod cost increases with heliozoan cell volume but this is cancelled out by the increased cell budget, giving short and long axopods similar relative construction costs (Figure 2E).

Costs of individual axopods and for heliozoan prey capture (the full set of axopods) were determined from structural data. These are compared (Figure 2F and 2G) to axopod and prey capture costs found to be optimal in our heliozoan cell division time calculations (see subsequent sections), revealing a rough agreement. Choanoflagellate prey capture costs (flagella + microvilli, Supplementary Information), as well as flagellar costs for a diverse set of species, were also included for comparison. Heliozoan prey capture is similar in cost to choanoflagellate prey capture and more expensive than flagellar construction cost, even for ciliates with 1000s of flagella.

### Heliozoan prey capture rate

Heliozoan prey capture rates were determined by simulation for two different modes of prey motility: diffusion and swimming. The model for the capture of diffusing prey was adapted from our previous work^58^ but was extended to different predator volumes and variation in axopod length. The model for the capture of swimming prey was established specifically for this work (Figure 3A-C). For both the diffusing and swimming prey models, prey capture rates were determined as a function of axopod length, axopod number, and predator volume (and prey volume for the swimming model). Capture rates are fitted with Equation 2 (Figure 3C), which is subsequently used in the cell division time calculations.

Prey capture rates can be expressed as clearance rates, i.e., external volume cleared of prey per unit time, and specific clearance rates, i.e., predator volumes cleared of prey per unit time. This reveals that simultaneously increasing axopod length and axopod number increases the capture rate by orders of magnitude (Figure 3D). Increasing axopod length, or number, individually has much more limited contributions to capture rate. A possible explanation is the following. The essential contribution to increased prey capture by axopods is that it creates an effective cell size much larger than that of the cell body. But if the axopod length is increased without an increase in the axopod number one can’t really speak of an effective sphere anymore. An increase in axopod number at fixed axopod length makes the cell increasingly approximate a sphere, but does not increase the effective size of the sphere. The clearance rate plot (Figure 3D) also reveals that the optimum axopod arrangement is far away from the highest clearance rate.

The prey capture rates found for the optima in the cell division time calculation (see subsequent sections) are converted into clearance rates and specific clearance rates and are compared to empirically determined values for a diverse set of species (Figure 3E and 3F). The heliozoan *Actinophrys sol*^43^ shows agreement with the clearance rates found in the swimming prey model, but not the diffusing prey model. Heliozoan clearance rates (including the swimming prey model) are at the higher end of the range found for a diverse set of protists and metazoa^79^. The diffusing prey model leads to clearance rates below the typical range, suggesting that it is not an effective strategy for nutrient acquisition. Taken together, the clearance rates for heliozoa, both empirical and from the model, show that axopod-based capture of swimming prey is highly effective but, in the light of Figure 2G, potentially more expensive than other mechanisms. For optimal axopod arrangements, prey capture locations were shown to favor the far ends of the axopods (Figure 3G-J), with possible implications for localization of adhesive-excreting extrusomes.

### Heliozoan cell division time and optimal axopod lengths and numbers

The cost of the prey capture mechanism (Eq. 1) adds to the amount of resources required for the duplication of the cell (Figure 4A, left panel), extending the cell division time. On the other hand, investment in a prey capture mechanism (Eq. 2) could increase the flux of resources into the cell (Figure 4A, middle panel), decreasing the cell division time. The combined effect of this on the cell division time is captured by Eq. 3 and leads to optimal axopod lengths and numbers at the minimal cell division time (Figure 4B). Fractional occupancy, *f_occ_*, of axopods on the cell body surface is significantly below 1 for the optima (Figure 4A, right panel, Supplementary Information).

**Figure 4:**
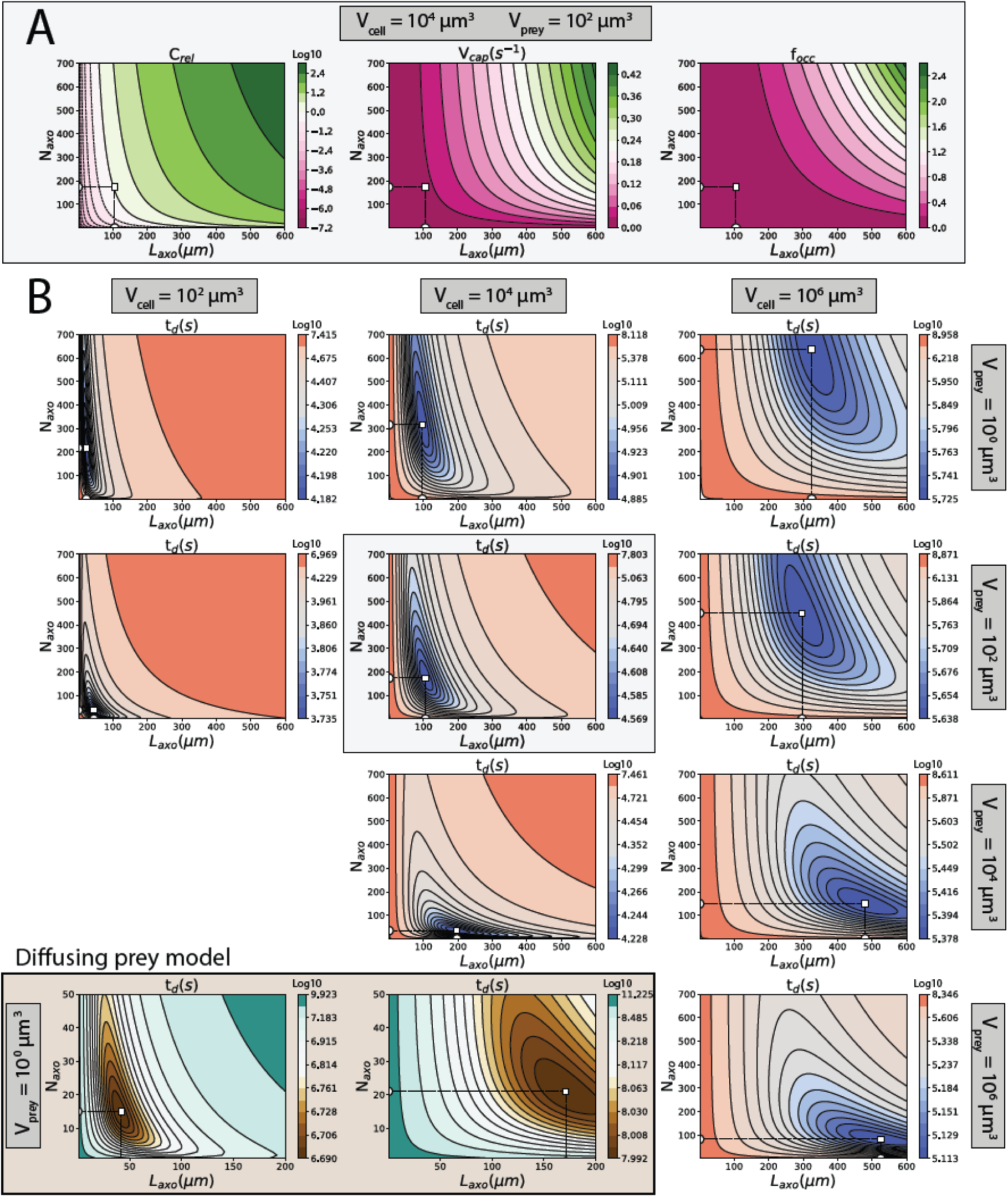
Cell division times and optimal axopod lengths and numbers. A) Relative cost (*C_rel_*), prey capture rate (*v_cap_*, at a *c_prey_* of 10^-8^ prey cells µm^-3^), and fractional cell-surface occupancy by axopods (*f_occ_*) as a function of axopod length and number. White square: position of minimal cell division time (Fig. 4B). B) Cell division times calculated with the swimming prey model and the diffusing prey model (lower left). White square: position of minimal cell division time. The contour levels for the cell division times were hyperbolically scaled to clearly visualize both the valley and steep slopes (Supplementary Information).

An increase in predator size leads to an increase in the optimum for both the axopod number and the axopod length, for both diffusing and swimming prey. An increase in prey volume, examined only for swimming prey, is associated with a steep decrease in axopod number, caused by the increased likelihood that larger prey collide with an axopod, and a more modest increase in axopod length. This has implications for understanding flagellate cell volume. An environment with an abundance of heliozoans would provide a selective pressure to reduce cell size for the flagellates because larger cells would experience a higher mortality. For the capture of diffusing prey, the optima lie at significantly lower axopod numbers than for swimming prey, courtesy of the cloud like nature of the diffusion paths compared to the straight paths of the swimmers.

The optimal axopod lengths, axopod numbers, and cell division times found in the models (Figure 4) can be compared to empirically determined values (Figure 5A–5C). Optimal axopod lengths for the diffusing and swimming prey, as well as for different prey sizes, are similar to each other and follow the same trend as the data, but are somewhat longer.

**Figure 5:**
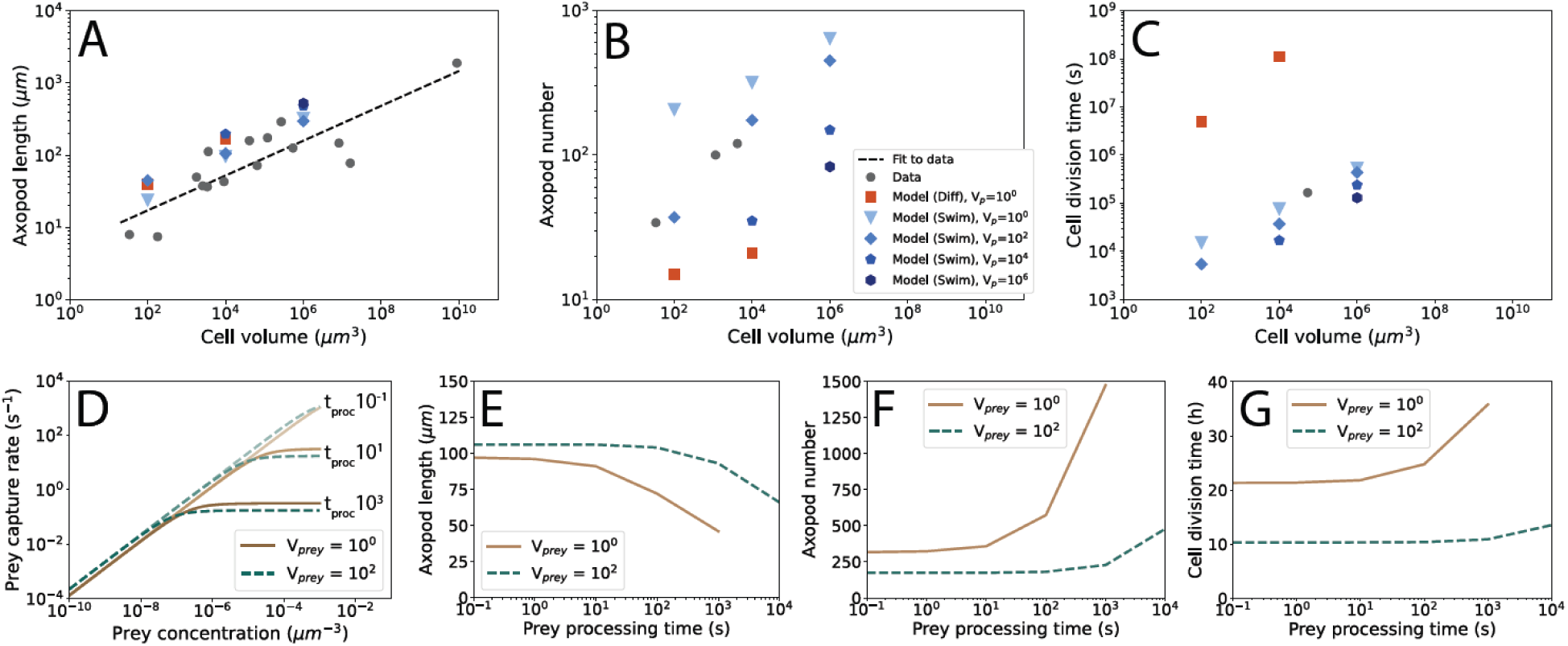
Comparing model results to experimental data and axopod saturation. A) Optimal axopod length compared to observations. Regression: *y* = 5.7*x*^0.24^. Legend in B. B) Optimal axopod number compared to observations. C) Minimal cell division time compared to observation. D-G) Results of the axopod saturation model. D) The prey capture rate as a function of prey concentration. Predator volume: 10^4^ µm^3^, prey volumes in µm^3^, axopod length: 106 µm, axopod number: 174, and prey processing times, *t_proc_*, ranging from 10^-1^ – 10^3^ seconds. E) Optimal axopod length, (F) optimal axopod number, and (G) minimal cell division time as a function of prey processing time.

There are only a handful of species of heliozoa with quantified axopod numbers and the comparison between model and data reveals a more complicated picture (Figure 5B). The predicted axopod numbers vary considerably but encompass the empirical values. Predictions for small *swimming* prey are higher than the observations and predictions for small *diffusing* prey are lower. As the swimming prey get larger, the optimal axopod number gets closer to the observed values, and then drops below it. Observed axopod numbers could result from intermediate prey sizes or prey size mixtures (as examined below).

For cell division times, there is likewise very sparse data, but the available datum falls within the range found for the swimming prey model (Figure 5C). The minimal cell division time found for heliozoans feeding on diffusing prey is orders of magnitude higher. This, in combination with the earlier findings on the clearance rates of diffusing prey (Figure 3E and 3F), suggests that feeding on small diffusing prey is a bad strategy.

### Axopod saturation

So far, the discussion has focused on the regime in which prey cells are too scarce to overwhelm the capacity of the prey capture mechanism. It is conceivable, however, that due to high local concentrations of prey, and/or long prey handling times, the axopods are occupied with a previous prey cell when a new prey presents itself. To account for this, a processing time, *t_proc_*, was introduced to calculate an effective axopod number (the number of axopods that are free to capture prey). This allows for the calculation of prey capture rate as a function of both prey concentration and prey processing time (Figure 5D), revealing that the prey capture rate saturates at a level determined by the prey processing time and that this happens at different prey concentrations.

The saturation model can be applied also to the question of cell division time minima and can be used to predict optimal axopod lengths and numbers as a function of prey processing time (Figure 5E and 5F). Optimal axopod length decreases after a threshold in prey processing time is reached (Figure 5E). The opposite happens with optimal axopod numbers, which increase as processing time increases (Figure 5F). Saturation of axopods decreases the rate at which prey is captured, increasing the cell division time (Figure 5G). Higher axopod numbers compensate for this, but come at a cost that also has a negative impact on growth. The decreased axopod length reduces the cost but also reduces the prey capture rate. The balance of all these considerations is that minimal cell division time increases with prey processing time.

### Evolutionary wandering of axopod length and number

Cellular properties vary over evolutionary time as has, for example, been observed in the extensive fossil record of diatoms^81^. The cellular properties of heliozoans must have varied over evolutionary time as well although the fossil record is too sparse to verify this^82^. Here, we examine the potential of two evolutionary forces to alter axopod length and number over time (Figure 6A): (1) genetic drift and (2) selection in response to long-term changes in prey composition.

**Figure 6:**
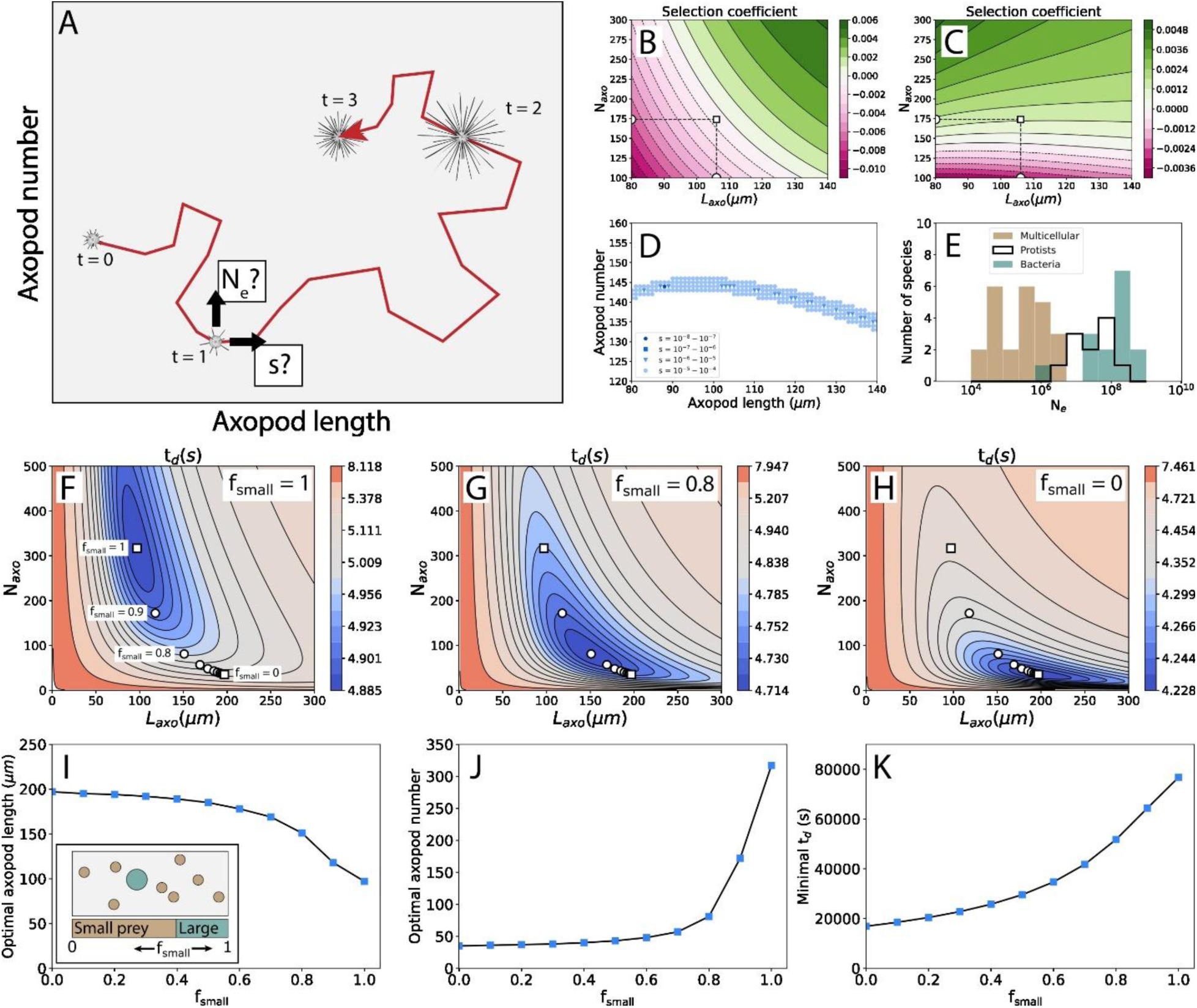
Evolution of axopod properties. A) Illustration of wandering axopod length and number over evolutionary time. B-E) Assessing the possibility of genetic drift changing the axopod length or number. B) The selection coefficient associated with changing the length of the axopods by 1 µm as a function of the starting axopod length and number. White square: position of minimal cell division time (Fig. 4B). C) Same as B but for changing the axopod number by one. D) Same as C but zoomed. E) Effective population sizes for various groups of organisms^80^. F-K) Effect of prey community composition on the optimal axopod length and number. F) Cell division times when the fraction of biomass (*f_small_*) belonging to small prey is 1. White points indicate optima for different fractions *f_small_* . G) Cell division times for *f_small_* = 0.8. H) Cell division times for *f_small_* = 0. I) Optimal axopod length as a function of small prey biomass fraction, *f_small_*. The inset illustrates the meaning of the biomass fraction, *f_small_*, with a 2D example. J) Optimal axopod number as a function of *f_small_* . K) Minimal cell division time as a function of *f_small_*.

Genetic drift has the potential to push axopod length and number to suboptimal values simply by the noise in the evolutionary process overpowering selection. The impact of genetic drift hinges on a comparison^83^ of the selection coefficient associated with a cellular change, *s*, and the effective population size, *N_e_*. Approximately, if *N_e_s* > 1 selection wins and can maintain the population close to the optimal (local) cellular arrangement, and if *N_e_ s* < 1 drift wins and the cellular arrangement can wander away from the optimum to some extent. We determined selection coefficients for plausible changes in the phenotype: a delta of one axopod in the axopod number and 1 µm in the axopod length. Selection coefficients were approximated using the normalized derivative of the cell division time (Eq. S29) and are shown for axopod length in Figure 6B and for axopod number in Figure 6C and 6D. For 1 µm changes in axopod length the selection coefficient is in almost all cases larger than 10^-4^. This is true also for addition or removal of single axopods, except that there is a thin band with a selection coefficient below 10^-4^ (Figure 6D). With effective population sizes showing a lower limit of 10^4^, it is unlikely that significant wandering of axopod length and number are caused by genetic drift. However, lower prey concentrations could cause tighter margins in the cost-benefit comparison and increase the influence of genetic drift.

An alternative source of evolutionary wandering of axopod length and number is variability in prey size. This is examined in a dual prey model (Eq. S30 and S31), in which the heliozoan can capture and consume prey of different cell volumes. Prey sizes are 1 and 10^4^ µm^3^ and their abundances are varied with the parameter *f_small_*, the volume fraction of small prey (with large prey having a volume fraction of 1 − *f_small_*). The total amount of prey volume, i.e. prey biomass, per unit of medium volume is constant. Cell division time is determined for ranges of axopod length and number, and for an *f_small_* between 0 and 1 (Figure 6F–6H). The minimal cell division time is found at different combinations of axopod length and number when *f_small_* is changed (Figure 6F–6K). A higher fraction of large prey also decreases the minimal cell division time, because larger cells are easier to capture. Remarkably, the larger prey has a significantly stronger effect than the smaller prey on the optimal axopod length and number (Figure 6I and 6J). With most of the change in optimal parameter values already having occurred when the fractional volume of the large prey is only 0.2 (a 4×10^4^-fold difference in cell numbers). Suggesting that direct observations of prey capture frequency could be highly misleading about prey impact on selective pressures.

## Discussion

### Explaining axopod length

Lengthening axopods causes increases in prey capture rate (Figure 3C, 3D, and 4A), and if axopod number rises as well, this increase appears to be indefinite. This is not reflected in an indefinite lowering of the cell division time as at relatively short axopod length a division time minimum is reached (Figure 4B). How can this be explained? The first step in the explanation is that increasing the axopod length also leads to an increase in the axopod cost. The axopod cost increases with the cube of the length, because of the observed linear relation between axopod length and width (Figure 2C). The increase in prey capture rate maximally scales with the square of the axopod length (due to the scaling of the surface area of a sphere). Typically, the increase in prey capture rate is even weaker, *γ* < 1.7 in Eq. 2. Thus, the cost will always outstrip the gain as axopods become longer causing a minimal cell division time at intermediate axopod length.

It remains to be explained why the width of the axopod scales linearly with its length. Axopods ought to be straight because the further each segment of an axopod is away from the cell the closer it approaches the ideal of an increase in effective cell radius. This is demonstrated by comparing the prey capture rates of pairs of heliozoan arrangements in which the total axopod length is equal, but in which one arrangement has two times the axopod length and the other two times the axopod number. These comparisons yield a 1.86±0.62 (mean±SD, N=107) higher capture rate for the arrangement with the longer axopods. Thus, an axopod is likely to be constructed in a way that keeps it straight. If forces on the axopod exceed a critical threshold, it will buckle. The force required for buckling, *F_crit_*, is given by (modified from^84^):

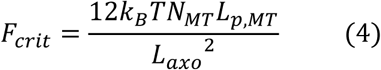

Here, *k_B_* is Boltzmann’s constant, *T* is temperature, *N_MT_* is the number of microtubules in an axopod, *L_p_*_,*MT*_ is the persistence length of an individual microtubule (a measure of the stiffness of the microtubule), and *L_axo_* is the axopod length. It is assumed in Eq. 4 that the persistence length of a bundle of microtubules is simply the sum of the persistence lengths of individual microtubules. If the typical forces that an axopod experiences are independent of its length, the number of microtubules should scale with the square of the axopod length, as is observed (Figure 2C). Optimal axopod length can thus (largely) be explained by a combination of the physics of the bending of rods and a biological cost-benefit tradeoff.

### Comparison with other filament-based cellular phenomena

There are parallels between the present axopod work and findings on other cellular filaments that suggest the possibility of a deeper understanding of cellular filaments. Receptors for signal detection have been found in the membranes of cilia. Like axopods, these cilia are slender filaments that extend away from the cell surface and like axopods these cilia must collide with particles, though in this case signal molecules not prey cells, and movement by diffusion not swimming. It has been found that a cilium can enhance detection (i.e. collision) of signaling molecules significantly over a patch of membrane with the same surface area as the cilium^85^. It was found that at the same length-to-width ratio found for axopods (∼190, Figure 2C) a patch on the cell surface must be ∼11-fold larger in area than the cilium to achieve the same detection sensitivity. A similar calculation for a heliozoan cell with one axopod finds that for a range of axopod lengths from 40-240 µm the fold difference in capture rate between axopod and surface patch, with the same area as the axopod, varies between 330-1 (*V_cell body_* = 10^4^ µm^3^, swimming prey).

Suctorians (ciliophora), like heliozoans, are festooned with cellular filaments, called tentacles. The tentacles differ from axopods in that prey is captured at the tips and prey cytoplasm is ingested through the center of the filament^23^. In recent work^37^ an analysis was performed on experimentally determined suctorian tentacle lengths and numbers that bears some similarity to the heliozoan model presented here. Their experimental findings were rationalized by a simple and provocative model that imagines a capture volume around each tentacle tip, with the capture volume bearing some relation to capture rate. At a critical tentacle number, the capture volumes start to overlap, leading to diminishing returns on investment. The simple model captures their empirical results neatly. A clear difference between the suctorian model and the heliozoan model presented here is that the total investment in tentacles isn’t calculated but simply observed (as a total tentacle length), whereas for the heliozoan model the optimal total investment is explicitly calculated.

### Simplifying the heliozoan prey capture model

Heliozoan prey capture rates were calculated by a simulation that includes an explicit heliozoan geometry and many *in silico* observations of prey capture events. A simpler analytical approach could be practically useful and provide insight into the prey capture mechanism. We examined three simple models for their ability to approximate the prey capture rates found with the full heliozoan prey capture model (Supplementary Information): (1) a model akin to the suctorian prey capture volume around a tentacle tip^37^, (2) a prey capture volume model that accounts for the axopod’s ability to capture prey along its entire length, and (3) an analytical calculation of prey capture probability that relies on a simplified (cube-shaped) axopod distribution. The resultant prey capture rates are divided by the heliozoan simulation-based capture rates (for all 199 swimming prey simulations) to obtain a ratio that yields the performance of the model relative to the full simulation: for model (1) the ratio ranges between 9.0×10^-4^-8.1 with a geometric mean of 4.7×10^-2^, for model (2) this is 0.0088-14 and 0.15, and for model (3) this is 0.18-1.4 and 0.61. The first model performs badly, in part due to representing the wrong prey capture architecture. The second model, that accounts for the axopod capture volume properly, works better but is still significantly off. The best performer is model three, which still has significant discrepancies with the full capture model, but which emphasizes the key importance of prey behavior in combination with the capture geometry in setting the capture rate.

## Conclusion

Prey capture mechanisms are some of the key cellular traits that phenotypically distinguish protist taxa. As such, understanding how the investment in prey capture mechanism relates to its effectiveness, and how the combined effects of cost and capture rate affect the cell division time, is crucial for understanding protist diversity. We have demonstrated that such an understanding is achievable for the heliozoan prey capture mechanism. The method presented here was applied to heliozoans first because the basic principles of their prey capture mechanism are relatively easy to model. There is no *a priori* reason why the methods applied would not work for other capture mechanisms, as long as a prey capture rate can be linked to an investment. Deep mechanistic understanding has been emerging for prey capture by flagellates, and this is a prime target for the application of the framework presented here.

## Acknowledgments

This work was funded by the Moore–Simons Project on the Origin of the Eukaryotic Cell, Simons Foundation 735927, the National Institutes of Health, R35-GM122566, and the National Science Foundation, DBI-2119963.

## Notes

### Competing Interest Statement

The authors have declared no competing interest.

