## Supplementary Information for "Heliozoan cellular architecture predicts prey capture performance and growth rate"

### Quantitative data collection on heliozoan morphology

All experimental data presented in this work were obtained from the literature, and include cell volumes, axopod widths, axopod lengths, microtubule density in the axopods, axopod number per cell, and cell division time (Table S1).

The dimensions of the heliozoan axopods are not systematically reported and, for this work, were in many cases estimated from published micrographs and figures. Some species were reported in the literature twice with different values and these were included as separate entries as they may represent genuine biological variation. Some of this data was included in our earlier work[1].

To estimate the cost of an axopod, estimates of the number of microtubules in an axopod are required. The number of microtubules per unit cross-sectional area was collected for three species of heliozoans (Table S1) giving an average of 819±177 $\mu m^{-2}$ (±SD). Combined with the width of an axopod, this density can be used to estimate microtubule costs of axopods. This data was included in our earlier work[1].

The number of axopods per heliozoan cell varies between species, as is clearly observed in micrographs, but its exact magnitude is frequently not reported and is difficult to estimate from published micrographs. Three estimates of heliozoan axopod number were obtained from the literature (Table S1). For *Heterophrys marina* the value was reported in the paper. For *Microheliella maris* the value was reported as >20 and we obtained an exact estimate from a drawing of the cell. Finally, for *Raphidiophrys contractilis* axopods were counted from a micrograph, which gives an underestimate because of the limited depth of focus. This number was extrapolated to the full cell number by assuming a focal depth of 3.5 μm, by using the cell surface area in the focused volume (approximated as a cylinder) and comparing that to the surface area calculated from the cell diameter.

**Table S1: Quantitation of heliozoan cellular morphology and ultrastructure.**

| Species | Cell body volume* ($\mu m^3$) | Axopod width (μm) | Axopod length (μm) | Microtubule density ($\mu m^{-2}$) | Axopod number | Ref. |
|---|---|---|---|---|---|---|
| ***Actinophrys salsuginosa*** | $1.2\times10^5$ | 1.642 | 175.0 | | | [2] |
| ***Actinophrys sol*** | $4.1\times10^4$ | 0.490 | 160.0 | | | [3] |
| ***Actinophrys sol*** | $6.5\times10^4$ | | 72.3 | | | [2] |
| ***Actinophrys tauryanini*** | $2.7\times10^5$ | 4.197 | 291.6 | | | [2] |
| ***Actinosphaerium eichhornii*** | $8.2\times10^6$ | 0.735 | 147.6 | | | [2] |
| ***Actinosphaerium nucleofilum*** | $1.6\times10^7$ | | 78.0 | 655 | | [2,4] |
| ***Berkeleyaesol magnus*** | $8.8\times10^9$ | | 1892.0 | | | [5] |
| ***Ciliophrys azurina*** | $3.4\times10^3$ | | 36.7 | | | [2] |
| ***Ciliophrys infusionum*** | $1.8\times10^2$ | | 7.5 | | | [6] |
| ***Ciliophrys infusionum*** | $2.6\times10^3$ | | 37.8 | | | [2] |

| | | | | | | |
|---|---|---|---|---|---|---|
| ***Ciliophrys marina*** | $1.8\times10^3$ | 0.100 | 50.0 | | | 7 |
| ***Clathrulina elegans*** | $9.1\times10^3$ | 0.193 | 43.3 | | | 8 |
| ***Dimorpha mutans*** | | | | 1006 | | 9 |
| ***Gymnosphaera albida*** | $5.4\times10^5$ | 1.250 | 126.6 | | | 10 |
| ***Heterophrys marina*** | $1.2\times10^3$ | | | | 100 | 11 |
| ***Microheliella maris*** | $3.4\times10^1$ | 0.092 | 8.0 | | 34 | 12 |
| ***Raphidiophrys contractilis*** | $3.6\times10^3$ | 0.293 | 112.7 | | 120 | 13 |
| ***Raphidiophrys elegans*** | | | | 796 | | 9 |

* This only includes the volume of the spherical cell body and excludes the axopods.

### Prey capture cost of heliozoans

To obtain cell division time we require both the rate of nutrient uptake and the total number of nutrients required. The total nutrient requirement is the sum of the nutrient requirement of the cell body and the axopods. Therefore, the cost of the axopods must be determined as a function of axopod length and number. The main structural features of axopods accounted for here are a bundle of microtubules from the center of the cell to the tip of the axopod and a membrane that covers the part of the axopod that extends away from the cell body. As axopods become longer, they need to contain more microtubules for structural support, therefore widening them and compounding the cost of both microtubules and membrane.

Axopod construction costs are calculated using a previously established framework for determining the energetic costs of basic cellular building blocks such as amino acids and lipids[14–16]. In this approach, costs are expressed in units of ATP hydrolyses but include opportunity costs. The opportunity costs are not physical ATP hydrolyses happening in the cell but parts of nutrient molecules that are used directly in biosynthesis. The cost in ATP of these building blocks then refers to the ATP that could have been retrieved from these molecules if they were used for energy generation.

Individual axopod costs are mathematically defined as:

$$C_{axo} = C_{mem} + C_{MT} \qquad (S1)$$

Here, $C_{mem}$ is the membrane cost given by:

$$C_{mem} = L_{axo} 2\pi r_{axo} C_{area} \qquad (S2)$$

and $C_{MT}$ is the microtubule cost given by:

$$C_{MT} = \frac{N_{MT} L_{MT} N_{pf} N_{AA,tub} C_{AA}}{d_{monomer}} \qquad (S3)$$

The axopod shape is approximated as a cylinder. Membrane costs are dependent on axopod length, $L_{axo}$, and the radius of the transverse axopod cross-section, $r_{axo}$, with the construction costs per μm$^2$ of membrane given by $C_{area}$ (with a value of 1.45×10$^9$ ATP μm$^{-2}$)[15,17]. Microtubule costs are dependent on the number of microtubules in the axopod, $N_{MT}$, and their lengths, $L_{MT} = r_{pred} + L_{axo}$, with $r_{pred}$ being the radius of the predator and $L_{axo}$ being the length of the axopod. Each microtubule consists of $N_{pf}$ protofilaments with the spacing between tubulin monomers in each protofilament given by $d_{monomer}$. Each monomer of tubulin contains a number of amino acids $N_{AA,tub}$, and each amino acid costs on average $C_{AA} = 29$ ATP.[14] The total absolute cost of prey capture, i.e. the cost of the full complement of axopods, in the models and for the examined species of heliozoans is then given by:

$$C_{cap} = C_{axo}N_{axo} \qquad (S4)$$

The number of axopods given by $N_{axo}$. For cell division time calculations, the axopod cost is calculated relative to the cost of the cell body, $\frac{C_{cap}}{C_{cell\ body}}$, giving a total relative cell cost of:

$$C_{rel,tot} = 1 + \frac{C_{cap}}{C_{cell\ body}} \qquad (S5)$$

With the cost of the cell body given by an empirical relation[14]:

$$C_{cell\ body} = 2.7 \times 10^{10} {V_{cell}}^{0.97} \qquad (S6)$$

With $V_{cell}$ being the volume of the cell body (excluding axopods) and only the costs of duplicating a cell being accounted for, i.e. no maintenance costs.

Axopod construction cost calculations for individual heliozoan species were calculated using either (1) an empirical relation between axopod length and axopod width (Fig. 2C) to obtain the missing widths from the available lengths (Fig. 2D and 2E) or (2) the specific lengths and widths determined for each species, yielding a more limited dataset (Fig. 2F and 2G). Due to lack of structural information, construction costs of centroplasts or axoplasts were not accounted for explicitly but are approximated by extending the axopod microtubules to the middle of the cell. The same holds for extrusomes on the axopods. The cost of the full complement of axopods on a single cell was determined by multiplying the individual axopod cost by the number of axopods. This was done for only two species, *Microheliella maris* and *Raphidiophrys contractilis*, because those were the only species for which all the required information, axopod number, width, and number, was available.

Axopod construction cost in the model uses the empirical relation between axopod length and width (Fig. 2C) to obtain the axopod width. Axopod length itself is varied in the model.

### Heliozoan prey capture rate – Diffusing prey

The heliozoan prey capture mechanism relies on collisions between the prey cells and the axopods. In the models it is assumed that once a prey hits an axopod, in any position along its length, it gets stuck. It is also assumed that the prey has no chance of escaping after it gets stuck and is transported down the axopod and engulfed by phagocytosis with perfect fidelity.

Prey can either diffuse or swim and both are considered in separate prey capture models. For the diffusing prey, Smoldyn simulations[18] are performed in which the heliozoan cell body is represented by a sphere and the axopods are cylinders. This is similar to an earlier heliozoan prey capture model by us[1]. Axopod positions are given by the Fibonacci method:

$$x_i = \mathrm{r}_{pred}\cos\left(\cos^{-1}\left(2\frac{i}{N}-1\right)-\frac{\pi}{2}\right)\cos\left(2\pi\frac{i}{\phi}\right) \qquad (S7a)$$

$$y_i = \mathrm{r}_{pred}\cos\left(\cos^{-1}\left(2\frac{i}{N}-1\right)-\frac{\pi}{2}\right)\sin\left(2\pi\frac{i}{\phi}\right) \qquad (S7b)$$

$$z_i = \mathrm{r}_{pred}\sin\left(\cos^{-1}\left(2\frac{i}{N}-1\right)-\frac{\pi}{2}\right) \qquad (S7c)$$

This gives a roughly equally spaced distribution of points over the surface of a sphere. Here, $\mathrm{r}_{pred}$ is the predator cell radius (excluding axopods). To obtain the coordinates for the tips of the axopods, the axopod length is added to the predator cell radius. $N$ is the total number of axopods, $i$ is the axopod index and $\phi$ is the golden ratio, $\frac{1+\sqrt{5}}{2}$.

Prey cells diffuse with a diffusion coefficient of 0.396 μm$^2$/s, calculated with the Stokes-Einstein equation:

$$D = \frac{k_B T}{6\pi\eta r_{prey}} \qquad (S8)$$

Using a prey radius $r_{prey}$ of 0.62035 μm (which gives a prey volume of 1 μm$^3$), a temperature T of 298 K, and a viscosity $\eta$ of 8.891×10$^{-4}$ Pa·s (viscosity of water at a temperature of 298 K).

In the Smoldyn simulation, axopod radii are set as a sum of the real axopod radius and the radius of the prey cell. The reason for this is that Smoldyn particles are points without a shape or size that represent the center point of the prey cell. Two heliozoan cell volumes are considered, 10$^2$ and 10$^4$ μm$^3$, and one prey volume, 1 μm$^3$. Axopod lengths and number were varied and for each combination a separate simulation was performed. Prey was generated at a fixed rate at a spherical boundary surface with a radius of 1100 μm and concentric with the heliozoan cell. This prey has three fates, (1) it can diffuse in the simulation volume for the entire duration of the simulation, (2) it can collide with the spherical boundary surface and be removed from the simulation, or (3) it can collide with an axopod and be counted as a capture event. Prey that collide with the heliozoan cell body bounce back and continue diffusing. The simulations were run for $2\times10^6$ seconds with a simulation timestep of 0.01 seconds. During the first $1\times10^6$ seconds, the simulation needed to equilibrate and only the second $1\times10^6$ seconds were used for analysis. Prey captures were tracked and divided by the capture period to calculate a prey capture rate. This capture data, as a function of axopod length ($L_{axo}$) and number ($N_{axo}$), is fitted with:

$$v_{cap} = \frac{A c_{prey} {L_{axo}}^{\gamma} N_{axo}}{B + N_{axo}} \qquad (S9)$$

Prey capture rate depends on the fit parameters, $A$, $B$, and $\gamma$, as well as on the prey concentration $c_{prey}$. The main purpose of this equation is to have an analytical expression of prey capture that can be used in the cell division time calculations at arbitrary values of $L_{axo}$ and $N_{axo}$. The simulation itself doesn't contain prey concentration explicitly, but this can be calculated using a separate set of simulations that examine the rate at which a sphere with a radius of $\mathrm{r}_{pred} + L_{axo}$ captures prey cells. This is accomplished by relating the flux at this boundary to the concentration using[19]:

$$J(r) = -\frac{D c_{prey} a}{r^2} = -\frac{D c_{prey}\left(\mathrm{r}_{pred} + L_{axo}\right)}{\left(\mathrm{r}_{pred} + L_{axo}\right)^2} \qquad (S10)$$

Where, $c_{prey}$ is the prey concentration far from the sphere, $D$ is the diffusion coefficient of the prey cell, $a$ is the absorbing boundary (equal to $\mathrm{r}_{pred} + L_{axo}$ in our case), and $r$ is the distance from the center at which the flux is examined (also equal to $\mathrm{r}_{pred} + L_{axo}$). Rewriting yields:

$$c_{prey} = \frac{J\left(\mathrm{r}_{pred} + L_{axo}\right)\left(\mathrm{r}_{pred} + L_{axo}\right)}{D} \qquad (S11)$$

The minus sign has been omitted because we are only interested in the magnitude of the flux, not its direction. The flux, $J\left(\mathrm{r}_{pred} + L_{axo}\right)$, meaning $J$ as a function of $\left(\mathrm{r}_{pred} + L_{axo}\right)$, is obtained from the simulations. The prey concentration from Eq. S11 can be substituted into Eq. S9 to find A. Once the values of A, B, and $\gamma$ have all been determined the prey capture rate can be computed at any combination of $L_{axo}$, $N_{axo}$, and $c_{prey}$.

### Heliozoan prey capture rate – Swimming prey

In the watery environments that heliozoans inhabit, many single-celled organisms and small animals swim. Because swimming is much faster and on a straighter trajectory than diffusion, this is likely to have consequences for heliozoan prey capture. To account for this prey behavior a second set of simulations was performed, using a custom Python script.

The prey capture rate of swimming prey was obtained by splitting the problem into two separate simulations. In the first simulation a prey capture rate is determined for a perfectly absorbing sphere of radius $\mathrm{r}_{pred} + L_{axo}$ (the sum of the radius of the predator cell body and the length of the axopods). In the second simulation, a probability is determined of a prey cell hitting one of the axopods once it is within the capture sphere from the first simulation. The product of this probability with the sphere prey capture rate gives the prey capture rate of the heliozoan. The reason for splitting the simulation in two is to speed up the simulation. Hitting an axopod requires a much shorter simulation timescale than hitting a sphere and separating the simulation allows for speeding up the calculations of all the prey movements outside of the radius $\mathrm{r}_{pred} + L_{axo}$ where interaction with axopods is impossible.

In the first simulation, the capture of prey by a sphere, prey positions are generated randomly within a space defined by two concentric spheres. The radius of the inner sphere is $\mathrm{r}_{pred} + L_{axo}$ and the radius of the outer sphere is $20(\mathrm{r}_{pred} + L_{axo})$. These positions are created by selecting from a uniform distribution in the x, y, and z directions between $-20(\mathrm{r}_{pred} + L_{axo})$ and $+20(\mathrm{r}_{pred} + L_{axo})$. Only positions that lie outside the inner sphere and inside the outer sphere, calculated using $r = \sqrt{x^2 + y^2 + z^2}$, are used in the simulations. The individual prey cells are then given a random swim vector by first picking two angles from a uniform distribution, a polar angle between 0 and π and an azimuthal angle between 0 and 2 π, and then using these angles to calculate the three elements of the swim vector:

$$x = r_{step} \sin(\theta) \cos(\varphi) \qquad (S12a)$$

$$y = r_{step} \sin(\theta) \sin(\varphi) \qquad (S12b)$$

$$z = r_{step} \cos(\theta) \qquad (S12c)$$

Here, $r_{step}$ is the distance traveled per simulation timestep. The value of $r_{step}$ is set by dividing the diameter of the inner sphere by 10, $r_{step} = \frac{2(\mathrm{r}_{pred}+L_{axo})}{10}$. The timestep, $t_{step}$, of the simulation is given by $\frac{2(\mathrm{r}_{pred}+L_{axo})}{10 v_{swim}}$, with $v_{swim}$ being the swimming speed (set to 50 μm $s^{-1}$). The simulation is run for $N_{steps} = \frac{t_{sim}}{t_{step}}$ number of steps, with $t_{sim}$ as the overall simulation time. At the start of the simulation 1000 prey are generated by the above-mentioned method. In each timestep, the positions of the prey are updated by adding the swim vector. If a prey hits the outer sphere, i.e. if its distance from the center exceeds $20(\mathrm{r}_{pred} + L_{axo})$, it is removed from the simulation, and a new prey is added in the manner described above. If the prey hits the inner sphere, i.e. if its distance from the center drops below $(\mathrm{r}_{pred} + L_{axo})$, it is counted as a capture event, is removed from the simulation and a new prey is added in the manner described above. During each timestep a prey has a probability of changing its swim vector in a random direction. This probability is given by $\frac{1}{N_{straight}}$, with $N_{straight}$ being the average number of timesteps that a prey swims straight:

$$N_{straight} = \frac{L_{swim}}{r_{step}} \qquad (S13)$$

The straight swim length, $L_{swim}$, is 500 µm, and $r_{step}$ is the distance travelled per simulation timestep.

The number of capture events is monitored over time, and the total number of captures is divided by the simulation time to obtain a preliminary prey capture rate, $v_{cap,pre}$. Total number of captures ranged between 646-23040 and simulation time varied between 10000-240000 s. Prey concentration in the simulations, $c_{prey,sim}$, is calculated by dividing the number of prey, 1000, by the volume of the space between the surfaces of the two concentric spheres, i.e. $V_{outer\ sphere} - V_{inner\ sphere}$. The final prey capture rate, $v_{cap}$, is calculated by normalizing to a prey concentration, $c_{prey}$, of $10^{-6}$ prey µm$^{-3}$, using $v_{cap} = \frac{c_{prey}}{c_{prey,sim}} v_{cap,pre}$.

In the second simulation, the probability of hitting an axopod is calculated for prey cells that have hit the inner sphere. There is no direct coupling between the two simulations. Prey cells in the second simulation are generated on the inner surface of a bounding sphere at random locations, the bounding sphere being equal in radius to the inner sphere in the first simulation. The starting coordinates of the prey cells are calculated from a polar, $\theta_{pos}$, and an azimuthal, $\varphi_{pos}$, angle. For the polar angle, first a random number was drawn from a uniform distribution between -1 and 1, and then this number was transformed into an angle by using $\cos^{-1}(number)$. For the azimuthal angle a random number was drawn from a uniform distribution between 0 and 2π. The prey coordinates are calculated from these angles using equation S12 but replacing $r_{step}$ with the radius of the bounding sphere, $r_{inner}$.

The range of possible swim vectors can be represented by an inwardly pointing hemisphere normal to the inner surface of the bounding spere at the starting position of the prey cell. To generate the swim vectors, we start with a base vector, $\boldsymbol{v_{start}} = \begin{pmatrix} 0 \\ 0 \\ -1 \end{pmatrix}$.

This can be pictured as a downward pointing arrow that is located at the top of the sphere. The direction of this arrow is modified using a polar angle, $\theta_{swim}$, and an azimuthal angle, $\varphi_{swim}$. First, the two angles are drawn from a distribution using:

$$\theta_{swim} = \frac{\cos^{-1}\left(1 - \text{Uniform}\left[\frac{\pi}{2}\right]\right)}{\cos^{-1}\left(\frac{\pi}{2}\right)} \frac{\pi}{2} \qquad (S14a)$$

$$\varphi_{swim} = \text{Uniform}[2\pi] \qquad (S14b)$$

Here, $\text{Uniform}[x]$ is sampling from a uniform distribution between 0 and $x$. This ensures that all directions are sampled uniformly, as a prey cell can come from any direction represented by a hemisphere. The base vector is then rotated along the y- and z-directions using the rotation matrices:

$$\boldsymbol{M_{swim,y}} = \begin{pmatrix} \cos(\theta_{swim}) & 0 & \sin(\theta_{swim}) \\ 0 & 1 & 0 \\ -\sin(\theta_{swim}) & 0 & \cos(\theta_{swim}) \end{pmatrix} \qquad (S15a)$$

$$\boldsymbol{M_{swim,z}} = \begin{pmatrix} \cos(\varphi_{swim}) & -\sin(\varphi_{swim}) & 0 \\ \sin(\varphi_{swim}) & \cos(\varphi_{swim}) & 0 \\ 0 & 0 & 1 \end{pmatrix} \qquad (S15b)$$

The dot product of $\boldsymbol{M_{swim,y}}$ and $\boldsymbol{v_{start}}$ yields a second vector, $\boldsymbol{v_2}$, which is used in the second dot product $\boldsymbol{M_{swim,z}} \cdot \boldsymbol{v_2}$ to obtain a third vector, $\boldsymbol{v_3}$. Next, the position of the vector needs to be moved to the starting coordinates of the prey. This is accomplished by another set of rotation matrices:

$$\boldsymbol{M_{pos,y}} = \begin{pmatrix} \cos(\theta_{pos}) & 0 & \sin(\theta_{pos}) \\ 0 & 1 & 0 \\ -\sin(\theta_{pos}) & 0 & \cos(\theta_{pos}) \end{pmatrix} \qquad (S16a)$$

$$\boldsymbol{M_{pos,z}} = \begin{pmatrix} \cos(\varphi_{pos}) & -\sin(\varphi_{pos}) & 0 \\ \sin(\varphi_{pos}) & \cos(\varphi_{pos}) & 0 \\ 0 & 0 & 1 \end{pmatrix} \qquad (S16b)$$

Here, $\theta_{pos}$ and $\varphi_{pos}$ are the angles that define the starting positions of the prey. These are used in another two rounds of dot products to produce the vector $\boldsymbol{v_5}$. The swim vector is given by:

$$\boldsymbol{v_{swim}} = \frac{2(r_{axo} + r_{prey})}{10} \boldsymbol{v_5} \qquad (S17)$$

Here, $r_{axo}$ is the radius of the axopod and $r_{prey}$ the radius of the prey cell. The denominator includes a 10 so that in the simulation the axopods are not missed because of steps that are too large.

In the simulation, prey cells are examined sequentially. A prey cell is moving step-by-step in a direction set by the starting position and the swim vector. If the prey cell crosses the bounding sphere the round is terminated and a new prey cell is initiated. If a prey cell hits an axopod it is counted as a capture, the round is terminated, and a new prey cell is initiated. This continues for $10^4$ rounds.

If a prey cell collides with the predator cell body, it attains a new swim vector. The new swim vector is generating like the initial swim vector but with a couple of changes. The starting vector is changed to:

$$\boldsymbol{v_{start}} = \begin{pmatrix} 0 \\ 0 \\ 1 \end{pmatrix} \qquad (S18)$$

The angles that define the new starting position are obtained from the coordinates of the collision site ($x$, $y$ and $z$):

$$\theta_{pos} = arctan2\left(\sqrt{x^2 + y^2}, z\right) \qquad (S19a)$$

$$\varphi_{pos} = arctan2(y, x) \qquad (S19b)$$

The equations for determining the swim direction, Eq. S14, and the rotation matrices, Eq. S15 and S16, remain the same.

A prey cell is considered to hit an axopod if it comes to a distance of $r_{axo} + r_{prey}$ from the center line of the axopod. In each timestep, the position of the prey cell is examined sequentially against the positions of all the axopods. The direction in which each axopod points, is defined by two angles:

$$\theta_{axo} = \cos^{-1}\left(2\frac{i}{N} - 1\right) - \frac{\pi}{2} \qquad (S20a)$$

$$\varphi_{axo} = 2\pi\frac{i}{\phi} \qquad (S20b)$$

Here, $N$ is the number of axopods, $i$ is the axopod index, and $\phi = \frac{1+\sqrt{5}}{2}$ is the golden ratio. The position of the prey cell is transformed by these two angles in such a way that the axopod being examined is pointing in the positive z-direction. This is accomplished by the dot product of the rotation matrices, Eq. S15 (with $-\theta_{axo}$ instead of $\theta_{swim}$ and $-\varphi_{axo}$ instead of $\varphi_{swim}$), and the prey position vector. If the prey cell in its new position ($x_{prey}$, $y_{prey}$, $z_{prey}$) satisfies both $z_{prey} > r_{pred}$ and $\sqrt{{x_{prey}}^2 + {y_{prey}}^2} < \left(r_{axo} + r_{prey}\right)$ it is considered to have collided with the axopod.

The probability of prey capture is then the number of all capture events divided by the total number of prey cells examined in the simulation. The prey capture rate is the product of the probability of prey capture and the prey capture rate by a sphere. Prey captures rates are determined for a range of axopod numbers and lengths with the range set by trial and error, making sure that the axopod number and length associated with the minimal cell division time (calculated later) fall within the ranges to avoid errors of extrapolation.

To be able to use the relationship between axopod number, axopod length, and prey capture rate effectively in the subsequent calculations, an equation is fitted to the simulation results. This is the same Eq. S9 as was used for the diffusing prey model.

### Calculation of the clearance rate

To facilitate comparison of the modeled heliozoan prey capture rates and empirically determined capture rates, the prey capture rate was converted into a clearance rate and a specific clearance rate. The clearance rate shows how much of the surrounding fluid is cleared of prey per unit time and the specific clearance rate expresses this per unit predator cell volume, i.e. $\frac{\text{clearance rate}}{V_{pred}}$. The clearance rate is given by:

$$\text{Clearance rate} = \frac{v_{cap}}{c_{prey}} \qquad (S21)$$

Here, $v_{cap}$ is the prey capture rate and $c_{prey}$ is the prey concentration. Combining this with Eq. S9 gives:

$$\text{Clearance rate} = \frac{A{L_{axo}}^{\gamma} N_{axo}}{B + N_{axo}} \qquad (S22)$$

### Prey capture positions

In both the diffusing prey and the swimming prey capture models, the coordinates of the prey capture sites are recorded during the simulation. These coordinates were used to compute a distance from the center of the cell using $d = \sqrt{x^2 + y^2 + z^2}$.

### Calculating the fractional cell-surface occupancy by axopods

To examine whether the surface of the cell body is not overcrowded with axopods the fractional cell-surface occupancy of axopods, $f_{occ}$, is determined. For this, the transverse cross-sectional area of an axopod is obtained from its radius which in turn is a function of the axopod length. This area is multiplied by the number of axopods to obtain the total area occupancy. To get a fraction, the area occupancy of axopods is divided by the area available on the surface of the cell body.

### Calculating cell division times

Combining the prey capture costs and the prey capture rate into one model allows for the calculation of the cell division time as a function of axopod number and length. The total amount of resources that the cell needs to gather to duplicate itself is the sum of the resources required for the cell body and the resources required for the axopods. The more axopods, and the longer the axopods, the more resources a cell needs to gather to duplicate, increasing cell division time. Increases in the number and lengths of axopods also increases the prey capture rate and thereby increases the flux of resources to the cell, reducing the cell division time. This is captured in the following equation:

$$t_d = \frac{\text{Nutrient requirement}}{\text{Nutrient uptake rate}} = \frac{N_{vol,pred}V_{pred}\left(1 + \frac{C_{cap}}{C_{cell\ body}}\right)}{f_{eff}N_{vol,prey}V_{prey}v_{cap}} \qquad (S23)$$

Here, $N_{vol,pred}$ and $N_{vol,prey}$ are the number of nutrients required for building the predator or prey cell per unit volume, $V_{pred}$ and $V_{prey}$ are the cell volumes of the predator and prey cells (for the predator the axopods are excluded from the cell volume), $C_{cap}$ is the absolute prey capture cost, $C_{cell\ body}$ is the cost of the predator cell body (excluding the axopods), $f_{eff}$ is the efficiency with which prey biomass is converted into predator biomass, and $v_{cap}$ is the prey capture rate. Both $C_{cap}$ and $v_{cap}$ depend on the number and lengths of the axopods (Eq. S1-S4 and S9).

### Saturation of the prey capture mechanism

In the previous sections it was assumed that the prey capture mechanism doesn't saturate. Saturation can be accounted for by including a processing time, $t_{proc}$, which is the time it takes an axopod to transport a prey cell down to the cell body and restore itself to a fully functional state (which may, for example, include recycling of extrusomes). The number of axopods that are occupied with prey is given by:

$$N_{axo,occ} = v_{cap}t_{proc} \qquad (S24)$$

This gives an effective axopod number of:

$$N_{axo,eff} = N_{axo} - N_{axo,occ} = N_{axo} - v_{cap}t_{proc} \qquad (S25)$$

The effective axopod number is associated with a reduced prey capture rate (modified from Eq. S9):

$$v_{cap} = \frac{Ac_{prey}{L_{axo}}^{\gamma}N_{axo,eff}}{B + N_{axo,eff}} \qquad (S26)$$

Combining Eq. 25 and 26 gives:

$$N_{axo,eff} = N_{axo} - \frac{Ac_{prey}L_{axo}{}^{\gamma}N_{axo,eff}}{B + N_{axo,eff}}t_{proc} \qquad (S27)$$

Rewriting this gives:

$$N_{axo,eff}{}^{2} + \left(B + AL_{axo}{}^{\gamma}c_{prey}t_{proc} - N_{axo}\right)N_{axo,eff} - BN_{axo} = 0 \qquad (S28)$$

$N_{axo,eff}$ can be obtained by using the quadratic formula, giving two answers. The sensible answer is plugged back into Eq. S26 to obtain the prey capture rate and to calculate an adjusted cell division time (Eq. S23). The minimal cell division time then gives the optimal axopod length and axopod number.

### Calculation of selection coefficients

By finding the minimal cell division time it is possible to obtain the optimal axopod length and number. Whether the optimal values are found in nature is not obvious from the fact that there is an optimum. Aside from the fact that not all cellular traits or environmental factors are accounted for, there is also the shallowness of the basin around the optimum to consider. If the difference in benefit between having N axopods or N+1 axopods is small, genetic drift may overcome selection, and the cell will typically not attain the optimum axopod number. A similar consideration holds for axopod length. To find out whether genetic drift or selection wins, we need to examine the selection coefficients associated with changes in axopod number and length and compare these with the effective population size. Selection coefficients can be calculated by taking the partial derivative of the cell division time with respect to either the axopod number or the axopod length. This is then normalized by dividing by the cell division time, giving the selection coefficients as:

$$s_{Laxo} = \frac{dt_d}{dL_{axo}}\frac{1}{t_d} \qquad (S29a)$$

$$s_{Naxo} = \frac{dt_d}{dN_{axo}}\frac{1}{t_d} \qquad (S29b)$$

These equations give the selection coefficient associated with a gain of length of 1 μm and a gain in axopod number of 1. Both equations are evaluated for a range of axopod lengths and axopod numbers.

### Dual prey model

Predators may target more than one type of prey for consumption. The model presented in the previous sections shows that prey size determines the optimal axopod length and number. Here, a dual prey model is introduced in which two prey, with different cell volumes, are present in the environment and can be captured and consumed by the same predator cell. The only thing that changes in a dual model is the nutrient acquisition rate, which is now dependent on two prey volumes and capture rates rather than one. Starting with Eq. S23 and including both a small and a large prey type we obtain:

$$t_d = \frac{N_{vol,pred} V_{pred} \left(1 + \frac{C_{cap}}{C_{cell\ body}}\right)}{f_{eff} N_{vol,prey} \left(V_{prey,small} v_{cap,small} + V_{prey,large} v_{cap,large}\right)} \qquad (S30)$$

To properly compare across different prey combinations, we introduce a fixed total prey volume per unit of medium volume, $c_{prey,vol}$. Eq. S9 is modified to include the prey biomass limit, yielding:

$$v_{cap,small} = \frac{A_{small} \left(f_{small} \frac{c_{prey,vol}}{V_{prey,small}}\right) L_{axo}{}^{\gamma_{small}} N_{axo}}{B_{small} + N_{axo}} \qquad (S31a)$$

$$v_{cap,large} = \frac{A_{large} \left((1 - f_{small}) \frac{c_{prey,vol}}{V_{prey,large}}\right) L_{axo}{}^{\gamma_{large}} N_{axo}}{B_{large} + N_{axo}} \qquad (S31b)$$

Here, $f_{small}$ is the volume fraction of the small prey compared to the total prey volume, and $V_{prey,small}$ and $V_{prey,large}$ are the cell volumes of large and small prey. This model can now be analyzed in the same way as the single prey model to find cell division times and the optimal axopod lengths and numbers at different relative abundances of prey types.

### Choanoflagellate prey capture cost

To put heliozoan prey capture costs in context, it is compared to choanoflagellate prey capture costs which consist of flagellum costs and microvilli costs. Stalks, that keep choanoflagellates stuck to a substrate, may also be relevant but are not included here. We examine four species of choanoflagellate for which enough information was available to perform a cost estimate: *Monosiga ovata*[20–22], *Salpingoeca amphoridium*[23], *Codosiga gracilis*[23], and *Stephanoeca diplocostata*[23]. Flagellum costs per unit length are obtained from Schavemaker and Lynch[17]. This is combined with the flagellum lengths from the four species of choanoflagellate to calculate the flagellum costs. The cost of the microvilli is approximated by using the cost per unit volume of a flagellum which, like a microvillus, is a cellular extension supported by cytoskeletal filaments and surrounded by a membrane. The volume of an individual microvillus is obtained from its cross-sectional area and its length, and the total volume of all microvilli on the cell is the individual microvillus volume multiplied by the microvilli number. The prey capture cost is the sum of the flagellum cost and the total microvilli cost and is expressed in units of ATP. A relative cost is calculated by dividing the prey capture cost by the cost of the cell body. The cost of the cell body is obtained by using the volume of the choanoflagellate cell body, excluding the flagellum and microvilli, and applying Eq. S6.

### Determining nutrient extraction efficiency

Not all nutrients present in the prey cells are converted into predator biomass. To account for this the fraction $f_{eff}$ was introduced which is the fraction of prey nutrients that is actually incorporated into the predator biomass and can have a value between 0 and 1. This number in not available for Heliozoans so an estimate was obtained from another protist, the ciliate *Balanion comatum*. Conversion efficiency of prey biomass into predator biomass was calculated by dividing the *Balanion comatum* cell volume by the total volume of prey *Rhodomonas* cells consumed during a cell cycle period, using data from Jakobsen and Hansen[24]. The comparison was carried out over a *Rhodomonas* concentration range of ~290-6900 cells $mL^{-1}$ , with a range of *Balanion* cell volumes of ~1300-2400 $\mu m^3$, and an average conversion efficiency of 0.29 was found. We used a value of ¼ in our calculations.

### Applying the suctorian capture volume model to heliozoan model results

A capture volume approach successfully captures some aspects of the tentacle number and length distribution in the suctorian *Podophrya collini*[25]. A similar calculation can be performed for heliozoan prey capture. The area of the bounding sphere concentric with the predator cell, and that goes around the axopod tips, was calculated using a radius equal to the sum of the predator cell body radius and the axopod length. Next, the cross-sectional area was determined for the prey cell. This cross-sectional area was multiplied by the optimal number of axopods to arrive at a measure similar, and for the present purpose equivalent, to the capture volume approach applied to the suctorian. This product was divided by the area of the bounding sphere to obtain a ratio. This ratio was multiplied by the sphere capture rate obtained in step one of the heliozoan prey capture simulations for swimming prey, yielding a prey capture rate. This calculation was performed for each combination of predator volume, prey volume, axopod number, and axopod length that were used in the heliozoan prey capture simulations for swimming prey ($N_{sim}$ = 199).

### Heliozoan prey capture rate based on fractional capture volume

For convenience in use, and for enhancing our mechanistic understanding, it would be helpful if simpler models could be found that reproduce the prey capture rates of the detailed heliozoan prey capture simulations. Similar to the tentacle capture volume in the suctorian work[25] a capture volume can be calculated specifically for axopods. This is defined, for each axopod, as the volume of a cylinder with a length equal to the axopod length a radius equal to the sum of the axopod cross-sectional radius and the radius of the prey. The total capture volume is the product of the cylinder volume and the number axopods. Next, the volume of the space around the predator between the tips of the axopods and the cell body is determined by determining the volume of the boundary spere and subtracting the volume of the predator cell body, $\frac{4\pi}{3}\left(r_{pred} + L_{axo}\right)^3 - \frac{4\pi}{3} r_{pred}{}^3$. The capture volume is then divided by this volume to obtain a capture fraction. This is multiplied with the sphere capture rate for swimming prey to obtain a capture rate. This calculation is performed for all parameter combinations used for swimming prey in the simulations ($N_{sim}$ = 199), axopod number, axopod length, predator volume, and prey volume, and compared to the prey capture rate observed in the simulation. Ratios of prey capture rates, $\frac{v_{cap,vol}}{v_{cap,sim}}$, give a range of 0.0088-14 with a geometric mean of 0.15.

### Prey capture rates from an axopod distribution in a cube

An issue with the fractional capture volume approach used above is that it does not account for the type of motility exhibited by the prey. Here, an approach is developed for the prey capture rate that accounts for the motility of the prey but also expresses the capture probability analytically. Just as with the swimming prey simulations this starts with the prey capture rate of a sphere, for which the simulations were used. The final capture rate is the product of the capture probability and the sphere capture rate.

The basic idea for obtaining an analytic estimate of the prey capture probability is to arrange the axopod length in a cube shaped space and launching prey cells at it in straight trajectories. To obtain the prey capture probability, for prey cells that have entered the bounding sphere, first a total axopod length is determined. This is separated into the three spatial dimensions by dividing by three to obtain the length of axopod available in each direction. For the next step the volume spanned by the axopods in the real heliozoan architecture is required, given by $\frac{4\pi}{3}\left(r_{pred} + L_{axo}\right)^3$. This is converted into a cube shaped volume with edges equal in length to the cube root of this volume. The length of axopod available to each dimension is divided by the edge length to obtain the number of axopod segments pointing in each direction. Finally, the distance between the axopod segments is determined by taking the square root of the number of axopod segments pointing into a single direction and using the resulting number to divide the cube edge length. The result is a cube shaped space crisscrossed in three directions by arrays of axopods (for simplicity the cell body is ignored in this calculation).

Next, the prey cells are considered to swim in straight paths at this axopod cube. They are assumed to swim only parallel to the x, y, and z axes. If that is the case, the probability of hitting an axopod is equal to an area fraction related to the square space between axopods, with sides the length of the axopod distance, and the radius of the prey cells. The prey capture rate is calculated by multiplying this probability with the rate of prey capture by a sphere (step 1 in the swimming prey simulation). Ratios of prey capture rates, $\frac{v_{cap,cube}}{v_{cap,sim}}$, give a range of 0.18-1.4 with a geometric mean of 0.61.

For two out of 199 calculations the space between axopods is so small compared to the prey size that we obtain a negative number. These were excluded from the analysis.

### Scaling of colors for the cell division time plots

To facilitate the visualization of both the small differences in the cell division time in the valleys and large differences on the steep slopes (Fig. 4) the contour levels are calculated from:

$$t_{d,contour,i} = \frac{b_1 y_i}{c_1 - y_i} + t_{d,minimal}$$

Here, $t_{d,contour,i}$ is the $i$'th contour level, $y_i$ is the $i$'th value picked at equal intervals between 0 and 0.97 (the number of intervals being equal to the number of contours minus one), $c_1$ is a constant set to 1, $b_1$ is a constant set to 0.1, and $t_{d,minimal}$ is the minimal cell division time in the plot.
